# XpressO-Melanoma: An Explainable Deep Learning Model for the Prediction of BRAF V600 Mutation Status in Cutaneous Melanomas

**DOI:** 10.1101/2025.07.03.662694

**Authors:** Vibha R. Rao, Thuy L. Phung, Shrey S. Sukhadia

**Affiliations:** Department of Pathology and Laboratory Medicine, Dartmouth Health, Lebanon, New Hampshire, USA; Department of Pathology and Laboratory Medicine, University of Texas Health Science Center at San Antonio, San Antonio, Texas, USA

**Keywords:** Cutaneous melanoma, BRAF V600E, whole-slide images, deep learning, explainable AI, computational pathology

## Abstract

*BRAF* V600E mutations are critical oncogenic drivers in cutaneous melanoma, influencing treatment decisions and outcomes. However, conventional molecular assays face limitations, including tissue availability, cost, and access. To address this, we present an explainable deep learning model that predicts *BRAF* V600E mutation status directly from diagnostic whole-slide images (WSIs) of skin cutaneous melanoma.

Using histopathological WSIs from The Cancer Genome Atlas (TCGA) and their corresponding mutation labels (BRAF wildtype vs. BRAF V600E), we trained a weakly supervised deep learning pipeline, XpressO, to identify tumor regions of interest (ROIs) predictive of BRAF mutation status. The model outputs attention heatmaps highlighting spatially relevant diagnostic features and computes a combined probability score from the top ten attention regions per WSI. These regions are further reviewed by a pathologist for biological appropriateness.

On an independent test set, the model achieved an AUC of 0.79 with balanced precision and recall, correctly identifying 7 of 8 BRAF V600E mutant cases. This demonstrates the model’s ability to capture phenotypic correlates of mutation status and highlights the potential of computational pathology in precision oncology. Our approach offers a scalable, interpretable, and cost-effective alternative to molecular testing, particularly in resource-limited settings.

## INTRODUCTION

Cutaneous melanoma is a highly aggressive malignancy of melanocytes (Siegel et al. 2019; Whiteman et al. 2016). Although it accounts for less than 5% of all skin cancers, it causes over 75% of skin cancer mortality, emphasizing its disproportionate clinical impact (Akbani et al. 2015). Increasing trends in melanoma incidence have been observed worldwide, emphasizing the growing clinical importance of early detection (Karimkhani et al. 2017). Molecular characterization of melanoma has revealed distinct genetic subsets, with activating mutations in the *BRAF* gene representing one of the most critical oncogenic drivers (Long et al. 2011). Approximately 40–60% of cutaneous melanomas harbor mutations in *BRAF*, predominantly the V600 variant, which results in constitutive activation of the *MAPK* signaling pathway, promoting tumor proliferation, survival, and metastasis (Davies et al. 2002; Flaherty et al. 2010). The identification of *BRAF* mutations has led to advances in targeted therapy, with selective *BRAF* and *MEK* inhibitors demonstrating significant improvements in progression-free and overall survival in metastatic melanoma patients (Ascierto et al. 2016; Robert et al. 2019).

Given the therapeutic implications, accurate determination of *BRAF* mutation status has become standard practice in the clinical management of advanced melanoma (Luke et al. 2017). Molecular assays such as allele-specific real-time polymerase chain reaction (RT-PCR), Sanger sequencing, and next-generation sequencing (NGS) are commonly employed to identify *BRAF* mutations (Cheng et al. 2018; Lee et al. 2011). Although sensitive and specific, these molecular techniques have inherent limitations, including dependency on adequate tumor cellularity, additional tissue consumption, longer turnaround times, and higher costs (Dummer et al. 2015). Immunohistochemistry (IHC) using mutation-specific antibodies such as VE1 offers a rapid and cost-effective alternative for detecting *BRAF* V600 mutations. However, IHC is intrinsically single-plex: the commonly used VE1 antibody detects only the BRAF p.V600E substitution on a single slide. Any additional biomarkers—or even non-V600E BRAF variants—require separate sections. This serial sectioning approach quickly exhausts tissue from shave or punch biopsies and inflates both cost and technologist time. Current reviews emphasize that this “one-marker-per-section” bottleneck is still the major limitation of routine IHC (Tan et al. 2020a). While emerging multiplex IHC/Immunofluorescence (IF) platforms can place 7–50 antibodies on a single slide, these techniques are for research-use-only and are not yet validated for *BRAF* mutation testing in clinical pathology labs, especially in the resource-limited regions (Harms et al. 2023; Tan et al. 2020b).

Recent advances in computational pathology and deep learning (DL) have enabled the extraction of actionable molecular information directly from diagnostic whole slide images (WSIs) (Bera et al. 2019; Echle et al. 2021; Sukhadia et al. 2025). Convolutional neural networks (CNNs) have shown strong ability to learn subtle histomorphologic patterns correlated with molecular alterations. In several cancer types, deep learning has been used to predict mutation status, subtype, or expression profiles directly from histopathology. Coudray et al. (Coudray et al. 2018) demonstrates that deep learning models can predict mutations in *EGFR, KRAS,* and *STK11* genes in lung adenocarcinomas directly from histopathology images. Similarly, Kather et al. (Kather et al. 2019) shows that microsatellite instability status could be accurately inferred from colorectal carcinoma WSIs using DL techniques. Expanding on this concept, Fu et al. (Y et al. 2020) applies DL to predict HER2 amplification from breast cancer histology slides, demonstrating the broader applicability of this strategy.

In melanoma, recent studies have demonstrated that *BRAF* mutation status can be predicted from whole slide images (WSIs) using deep learning approaches. CNNs have shown the ability to distinguish *BRAF*-mutant from BRAF wild-type melanomas based on histopathological features (O et al. 2022). Furthermore, advances in weakly supervised learning techniques, including attention-based multiple instance learning (MIL) frameworks, have enhanced the capacity of models to localize predictive tumor regions within WSIs without the need for pixel-level annotations (Cheng et al. 2025; Kim et al. 2022).

Despite promising performance, a critical limitation of many deep learning models remains their lack of interpretability, often referred to as the “black box” problem (Ghassemi et al. 2021). In medical domains, particularly pathology, explainability is essential to building clinician trust, ensuring diagnostic safety, and facilitating regulatory acceptance (A et al.; E and C). Models that can provide human-understandable rationales for their predictions are significantly more likely to be integrated into clinical workflows. Various strategies have been developed to enhance model transparency in computational pathology, aiming to link prediction outcomes with recognizable histological features (E and C 2020; Rj et al. 2019). Explainable deep learning approaches not only increase end-user confidence but also offer opportunities to uncover novel morphologic correlates of underlying molecular alterations, thereby bridging the gap between traditional histopathology and molecular diagnostics (D and S 2018).

Additionally, addressing spatial heterogeneity at the whole-slide image level remains a critical challenge in computational pathology. Melanomas often display complex architecture with intermixed tumor nests, inflammatory infiltrates, necrosis, and stromal components, posing challenges for consistent mutation prediction (Christie-Nguyen et al. 2025). Techniques such as patch-level attention weighting and region-based aggregation have been employed to focus model predictions on the most diagnostically relevant areas (Cheng et al. 2025; My et al. 2021). By selectively attending to tumor-rich and morphologically informative regions, models can improve both accuracy and biological relevance of predictions.

In light of these advances, we developed an explainable deep learning model “XpressO-melanoma” to predict *BRAF* V600E mutation status directly from diagnostic (Dx) WSIs of cutaneous melanoma. Leveraging curated cases from The Cancer Genome Atlas (TCGA), we repurposed our previously published DL pipeline called “XpressO” that performs tumor segmentation, feature extraction and classification of gene-expression labels, coupled with visualization of attention heatmaps of gene-expression on tissue slides, for prediction of *BRAF* V600E mutation labels. Our hypothesis is that deep learning models can identify morphologic patterns linked to BRAF-driven oncogenesis and use these patterns to predict mutation status, and that visual explanations of the model’s focus regions for that prediction will facilitate clinical trust and potential integration into diagnostic practice. This approach seeks to complement conventional IHC and molecular assays, particularly in scenarios with limited tissue availability or resource constraints, and represents a step toward spatially informed, AI-driven precision oncology in dermatopathology.

## RESULTS

### Model Evaluation of *BRAF* V600E Mutation Status

The XpressO-melanoma model predicted *BRAF* V600E mutation status from diagnostic WSIs of cutaneous melanoma. On the independent test set comprising 19 cases (11BRAF V600E wild-type (BVW) and 8 BRAF V600E mutant (BVE) melanomas, the model achieved an area under the receiver operating characteristic curve (AUC) of 0.79 (confidence interval: 0.57–1.00). The model’s precision, recall and F1-score were 0.72, 0.68, and 0.67, respectively. While the AUC of 0.79 suggests that the model effectively distinguished between BVW and BVE status based on morphologic patterns in WSIs, the precision and recall rates indicate that the model maintained a moderate level of predictive reliability while minimizing false positives and false negatives. The output of the XpressO-melanoma model (i.e., predictions of BVW versus BVE status for each WSI in the Test set) was inspected further using attention heatmaps as described in the following section.

**Figure 1:**
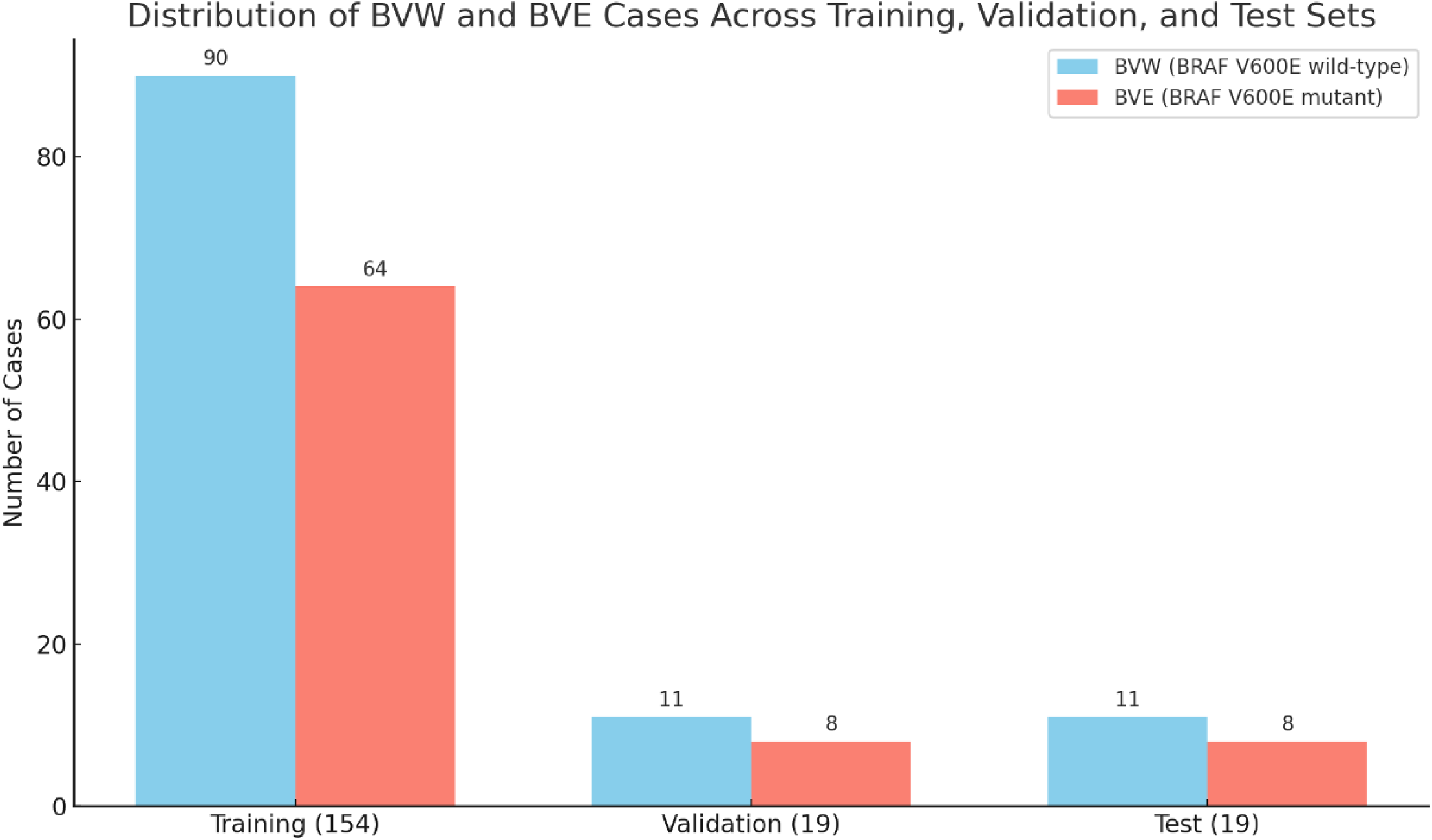
Distribution of BVW and BVE cases across Training, Validation, and Test Sets. Bar plot showing the number of BVW (blue) and BVE (red) cases across the training (n = 154), validation (n = 19), and test (n = 19) sets.

### Prediction and Visualization of Mutation-Associated Regions on WSIs

Attention heatmaps generated by the Heatmap module of the XpressO-Melanoma pipeline provided insights into the spatial localization of histological features associated with *BRAF* V600E mutation status. For each WSI in the test set, the attention module assigned scores to individual tissue patches, reflecting their relative contribution to the final slide-level prediction. The attention scores for the top 10 ranked patches were aggregated and mapped back to the original tissue sections, producing heatmaps that visually highlighted tumor regions driving the model’s prediction. Higher attention weights corresponded to areas shaded red on the heatmaps, while lower-weighted regions appeared with less intensity(blue).

Evaluation of heatmaps alongside the original slide-images revealed that the model correctly classified 87.5% (7 out of 8) *BRAF*-mutant cases, but it only correctly classified 54.54% (6 out of 11) *BRAF* wild-type cases (Table 1; Figures 2-5). Misclassifications primarily occurred near the decision threshold, where predicted probabilities for the incorrect class were relatively close to 0.5 in 4 out of total 19 cases (BVW+BVE) (Cases 5,6,9 and 11), leaving only 2 cases that were actual false positives (Cases 3 and 13) (Table 1).

**Figure 2:**
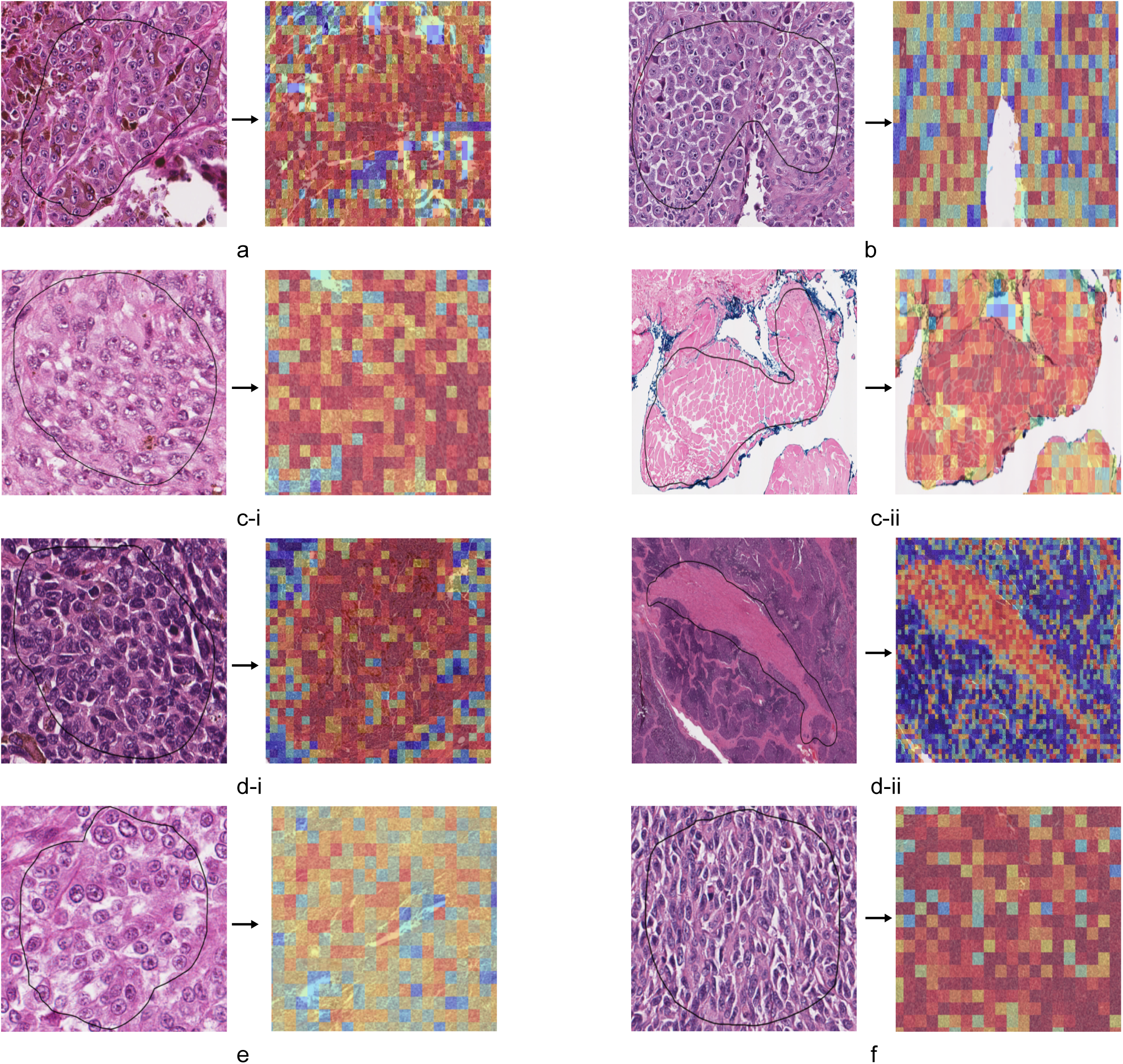
Representative Cases from Interpretation Category 1: BVW correctly predicted as BVW. **(a)** Case 1 -TCGA-EE -A182: Heatmap correctly predicted 83.5% probability for BVW and 16.4% probability for BVE. ROI outlined in black, **(b)** Case 2-TCGA-GN-A267: Heatmap correctly predicted 75.4% probability for BVW and 24.6% probability for BVE. ROI outlined in black, **(c-i)** Case 4-TCGA-EE-A29D: Heatmap correctly predicted 52.2% probability for BVW and 47.8% probability for BVE; heatmap shows attention over tumor areas. ROI outlined in black. **(c-ii)** Case 4-TCGA-EE-A29D:Heatmap correctly predicted 52.2% probability for BVW and 47.8% probability for BVE; heatmap shows attention over non-tumor areas. ROI outlined in black,**(d-i)** Case 7-TCGA-ER-A42H: Heatmap correctly predicted 57.9% probability for BVW and 42.1% probability for BVE; heatmap shows attention over tumor areas. ROI outlined in black,**(d-ii)** Case 7-TCGA-ER-A42H Heatmap correctly predicted 57.9% probability for BVW and 42.1% probability for BVE; heatmap shows attention over non-tumor areas. ROI outlined in black, **(e)** Case 8-TCGA-EE-A2MD: Heatmap correctly predicted 57.1% probability for BVW and 42.9% probability for BVE. ROI outlined in black, **(f)** Case 10-TCGA-GN-A264: Heatmap correctly predicted 58.6% probability for BVW and 41.4% probability for BVE. ROI outlined in black.

**Table 1:** Predicted BRAF V600E Mutation Status and Probabilities from Top-10 and Bottom-10Attended Patches, with Corresponding *BRAF, NRAS*, and *NF1* Mutation Statuses in the Independent Test Set.

| Case No. | TCGA_ID | Original label | Fig No. | Predicted mutation status using top-10 attended patches | Probability of mutation status using top-10 attention | Predicted mutation status using bottom-10 attended patches | Probability of mutation status using bottom-10 attention | *BRAF* | *RAS* | *NF1* |
| --- | --- | --- | --- | --- | --- | --- | --- | --- | --- | --- |
| 1 | TCGA-EE-A182 | BVW | 2 | BVW | 0.83535135 | BVE | 0.16464865 | - | - | +<br>*NF1*<br>*R440*<br>* |
| 2 | TCGA-GN-A267 | BVW | 3 | BVW | 0.75375223 | BVE | 0.24624783 | - | +<br>*NRAS*<br>*Q61*<br>*K* | - |
| 3 | TCGA-D3-A2J8 | BVW | 4 | BVE | 0.59119654 | BVW | 0.4088035 | - | +<br>*NRAS*<br>*Q61L* | - |
| 4 | TCGA-EE-A29D | BVW | 5 | BVW | 0.5215086 | BVE | 0.47849146 | - | - | - |
| 5 | TCGA-GN-A266 | BVW | 6 | BVE | 0.57945 | BVW | 0.42055 | - | - | - |
| 6 | TCGA-EB-A5SF | BVW | 7 | BVE | 0.50616527 | BVW | 0.49383473 | - | - | - |
| 7 | TCGA-ER-A42H | BVW | 8 | BVW | 0.57937396 | BVE | 0.42062604 | - | - | - |
| 8 | TCGA-EE-A2MD | BVW | 9 | BVW | 0.57118636 | BVE | 0.42881364 | - | +<br>*NRAS*<br>*Q61*<br>*R* | - |
| 9 | TCGA-EE-A2GR | BVW | 10 | BVE | 0.7521731 | BVW | 0.2478269 | - | - | +NF<br>1<br>C383<br>Yfs*1<br>0,<br>NF1<br>X258<br>0_spl<br>ice |
| 10 | TCGA-GN-A264 | BVW | 11 | BVW | 0.5860901 | BVE | 0.4139099 | - | - | - |
| 11 | TCGA-EE-A29X | BVW | 12 | BVE | 0.5390571 | BVW | 0.46094295 | - | +NR<br>AS<br>Q61<br>R | - |
| 12 | TCGA-D3-A2JN | BVE | 13 | BVE | 0.920806 | BVW | 0.07919398 | +BRA<br>F<br>V640E | - | - |
| 13 | TCGA-ER-A19C | BVE | 14 | BVW | 0.96872544 | BVE | 0.03127459<br>4 | +BRA<br>F<br>V640E | - | - |
| 14 | TCGA-D3-A1Q9 | BVE | 15 | BVE | 0.89735985 | BVW | 0.10264011 | +BRA<br>F<br>V640E | - | - |
| 15 | TCGA-ER-A2ND | BVE | 16 | BVE | 0.8608054 | BVW | 0.13919458 | +BRA<br>F<br>V640E | - | - |
| 16 | TCGA-D3-A2J7 | BVE | 17 | BVE | 0.8337906 | BVW | 0.16620935 | +BRA<br>F<br>V640E | - | - |
| 17 | TCGA-EE-A2MH | BVE | 18 | BVE | 0.7746738 | BVW | 0.22532617 | +BRA<br>F<br>V640E | - | - |
| 18 | TCGA-EE-A2GS | BVE | 19 | BVE | 0.58899146 | BVW | 0.41100854 | +BRA<br>F<br>V640E | - | - |
| 19 | TCGA-EE-A3AF | BVE | 20 | BVE | 0.75411 | BVW | 0.24588996 | +BRA<br>F<br>V640E | - | - |

### Interpretation Category 1: BVW correctly predicted as BVW

In Figure 2a (Case 1), the model correctly predicted the tumor as BVW with high confidence (83.5% probability). High attention (red) was assigned to tumor regions that aligned closely with hand-annotated areas, demonstrating accurate spatial localization. These regions exhibited ovoid-shaped nuclei, dense fibrous stroma, and well-circumscribed tumor boundaries which are morphologic features commonly associated with *BRAF* wild-type melanomas (De et al. 2020). This aligns with literature describing *BRAF*-wild-type tumors as having less nuclear pleomorphism and lower cellularity compared to *BRAF*-mutant melanomas, which tend to show more pronounced cytologic atypia (Kim et al. 2022). Molecular data from TCGA revealed the presence of an *NF1* mutation (R440*) and absence of *BRAF* or RAS mutations. *NF1*-mutant melanomas, especially desmoplastic subtypes, often contain tumor cells set in a collagen-rich matrix as seen in this case (T et al. 2015). This morphology likely contributed to the model’s strong confidence, as the absence of *BRAF*-driven cytologic hallmarks reinforced the *BRAF* V600E wild-type classification.

In Figure 2b (Case 2), the model correctly predicted a *BRAF* V600E wild-type status with a probability of 75.4%. The attention map revealed high attention (red) over tumor regions that aligned well with manually annotated areas, while non-tumor regions, including the epidermis, received low attention (blue), indicating effective spatial discrimination. Molecular profiling identified a pathogenic *NRAS* Q61K mutation, with wild-type *BRAF* and *NF1*. *NRAS*-mutant melanomas are typically more cellular and mitotically active than *NF1*-mutant or triple wild-type tumors but do not show the pronounced pleomorphism and epithelioid features seen in *BRAF* V600-mutant cases (B et al. 2011). These intermediate features may have contributed to the slightly lower confidence (compared to Case 1) in the model’s BVW prediction, as some areas may have resembled *BRAF*-mutant morphology without fully matching its hallmark traits. Nonetheless, the absence of hallmark *BRAF*-mutant features supported the accurate BVW classification.

In Figure 2c (Case 4), the model correctly predicted BVW status with a marginal probability of 52.2% for BVW and 47.8% for BVE mutation, indicating a high degree of uncertainty. The attention map revealed strong activation not only over tumor regions (Figure 2c-i) but also over normal skeletal muscle areas (Figure 2c-ii). This lack of clear separation may have contributed to the low confidence. Molecular profiling confirmed a triple-wildtype status, with no detectable mutations in *BRAF*, *NRAS*, or *NF1*. Triple-wildtype melanomas are known to exhibit variable and less distinct histologic patterns, which may overlap with both BVW and BVE features, making them particularly challenging for the model to classify (Smoller 2006). The attention to non-specific regions suggests that, in the absence of clear cytologic signals, the model may have relied on surrounding architectural or stromal cues, leading to the equivocal output.

In Figure 2d (Case 7), the model correctly predicted BVW status with 57.9% probability versus 42.1% for BVE, though with modest confidence. The heatmap revealed attention applied not only to tumor regions (Figure 2d-i) but also to surrounding eosinophilic stromal areas (Figure 2d-ii), suggesting some difficulty in distinguishing morphologically relevant features. The tumor was triple-wildtype, lacking mutations in *BRAF*, *NRAS*, and *NF1*. Such cases are often characterized by non-specific or overlapping histologic features, which may resemble both *BRAF* mutant and *BRAF* wild-type patterns (Kaczmarzyk et al. 2024). The modest prediction margin and non-specific attention suggest that the model may have been influenced by stromal elements or pale-staining regions rather than strictly tumor-specific morphology.

In Figure 2-e (Case 8), the model correctly predicted BVW with a probability of 57.1%, though with moderate confidence. The heatmap demonstrated moderate attention across most tumor regions on the H&E slide, indicating that the model effectively localized malignant areas. However, some regions contained a mixture of tumor and normal cells, which may have contributed to the model’s uncertainty. Molecular profiling revealed an *NRAS* Q61R mutation, with wild-type *BRAF* and *NF1*, indicating an *NRAS*-driven melanoma. *NRAS*-mutant melanomas often exhibit intermediate morphologic features which are more cellular than *NF1*-driven or triple-wildtype tumors but lacking the pleomorphism typical of *BRAF*-mutants (B et al. 2011). This may explain the moderate prediction confidence, as the model likely recognized increased cellularity without the hallmark features of *BRAF* V600E-mutant cytology.

In Figure 2-f (Case 10), the model correctly predicted BVW status with 58.6% probability. The attention map showed appropriate focus on tumor regions, which appeared predominantly spindle-shaped, features commonly associated with *BRAF*-wild-type melanomas(Kim et al. 2012). Despite the borderline confidence score, the model’s attention aligned well with tumor architecture, avoiding non-tumor areas. Molecular data confirmed a triple-wildtype profile (*BRAF*, *RAS,* and *NF1* wild-type), a subtype known to display histologic heterogeneity with less distinctive morphologic cues (Kaczmarzyk et al. 2024). The modest confidence score may reflect this ambiguity, but the overall prediction remained accurate.

### Interpretation Category 2: BVW incorrectly predicted as BVE

In Figure 3a (Case 3), the model showed uncertainty, assigning a 59% probability for BVE and 40% for BVW. Figure 3a-i displays tumor regions where the heatmap exhibited low attention (blue), despite the presence of atypical tumor cells with increased cellular density. These features are more consistent with a BRAF wild-type morphology (C et al. 2024), yet may have contributed subtle cues that partially resembled BRAF-mutant histology. Figure 3a-ii, by contrast, highlights necrotic regions that received disproportionately high attention (red) from the model. This misfocusing suggests that the model may have been influenced by non-specific necrotic and stromal features in the absence of clear cytologic hallmarks. Molecular data revealed the presence of an *NRAS* Q61L mutation, with no alterations in *BRAF* or *NF1*. Given that *NRAS*-mutant melanomas often show higher cellularity and proliferation than triple wild-type or *NF1*-driven tumors, but lack the cytologic hallmarks of *BRAF* V600E mutants, this case likely confused the model by presenting intermediate histologic cues (Kim et al. 2012).

**Figure 3:**
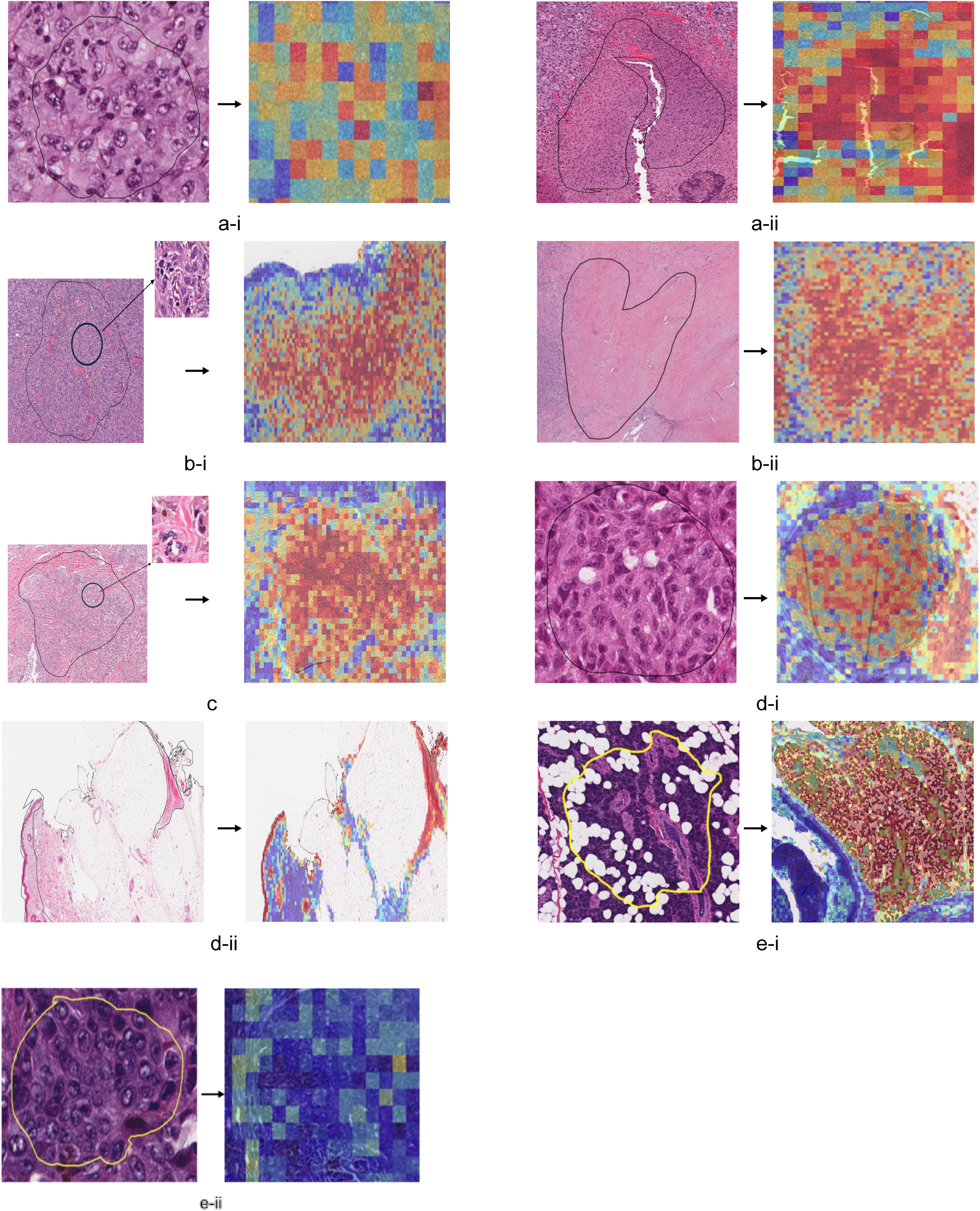
Representative Cases from Interpretation Category 2: BVW wild-type incorrectly predicted as BVE. **(a-i)** Case 3 – TCGA-D3-A2J8: Heatmap was equivocal with 59% probability for BVE versus 40% probability for BVW; heatmap shows low attention over tumor areas. ROI outlined in black,**(a-ii)** Case 3 – TCGA-D3-A2J8: Heatmap was equivocal with 59% probability for BVE versus 40% probability for BVW; heatmap shows high attention over non-tumor areas. ROI outlined in black,**(b-i)** Case5 - TCGA-GN-A266: Heatmap was inconclusive, assigning 58% probability for BVE versus 42% for BVW; heatmap shows attention over tumor areas. ROI outlined in black, **(b-ii)** Case 5 – TCGA-GN-A266: Heatmap was inconclusive, assigning 58% probability for BVE versus 42% for BVW; heatmap shows attention over non-tumor areas. ROI outlined in black,**(c-i)** Case 9 – TCGA-EE-A2GR: Heatmap falsely said 79% probability for BVE and 25% probability for BVW; heatmap shows attention over tumor areas. ROI outlined in black,**(c-ii)** Case 9 – TCGA-EE-A2GR: Heatmap falsely said 79% probability for BVE and 25% probability for BVW; heatmap shows attention over non-tumor areas. ROI outlined in black,**(d-i)** Case 11 – TCGA-EE-A29X: Heatmap was closely split with 54% for BVE and 46% for BVW; heatmap shows high attention over non-tumor areas. ROI outlined in yellow, **(d-ii)** Case 11 – TCGA-EE-A29X: Heatmap was closely split with 54% for BVE and 46% for BVW; heatmap shows low attention over tumor areas. ROI outlined in yellow.

In Figure 3b (Case 5), the model returned an ambiguous prediction, assigning 58% probability for BVE and 42% for BVW. The attention map highlighted both tumor cells (Figure 3b-i) and adjacent necrotic area (Figure 3b-ii). These regions contain dead cells that may have introduced noise that the model misinterpreted as suggestive of BRAF-mutant histology. Molecular profiling confirmed a triple-wildtype genotype, with no alterations in *BRAF*, *NRAS*, or *NF1*. Given the histologic ambiguity and stromal overlap common in triple-wildtype melanomas, the model’s misclassification reflects the broader challenge of distinguishing subtle morphologic cues in these cases (Kaczmarzyk et al. 2024).

In Figure 3c (Case 6), the model showed near-equal probabilities: 51% for BVE and 49% for BVW, reflecting high uncertainty. The heatmap revealed attention distributed across both tumor and non-tumor regions, including areas with spindle cells and fibrotic stroma. These overlapping histologic features may have contributed to the ambiguous prediction. Literature suggests that in cases with intermixed spindle cells and dense tumor nests, the model may struggle to discern *BRAF* wild-type from *BRAF* mutant features, particularly in areas of reactive stroma (A et al. 2008). The tumor was found to be triple-wildtype, lacking mutations in *BRAF*, *NRAS*, or *NF1*, which is consistent with reports that triple-wildtype melanomas often display heterogeneous and less distinct morphology that can mimic both BVW and BVE patterns (Kaczmarzyk et al. 2024). In such cases, the model may have responded to reactive or dense stromal components rather than clear-cut cytologic cues, underscoring the challenges of distinguishing mutation status in morphologically ambiguous slides.

In Figure 3d (Case 9), the model incorrectly predicted BVE with a high probability of 79%, despite the tumor being wild-type for *BRAF*. The attention map showed high attention (red) not only on tumor regions (Figure 3d-i) but also on non-tumor areas, particularly the epidermis and surrounding collagen-rich zones (Figure 3d-ii). This misclassification suggests that the model may have been influenced by non-specific architectural features or reactive tissue elements, mistaking them for *BRAF*-associated morphology. Molecular profiling confirmed *NF1* mutations (C383Yfs*10 and X2580_splice), with wild-type *BRAF* and RAS. *NF1*-driven melanomas typically lack the prominent pleomorphism and epithelioid cytology of *BRAF* V600E-mutants, which the model failed to account for here, likely due to misleading attention patterns over histologically ambiguous regions (B et al. 2011).

In Figure 3e (Case 11), the model incorrectly predicted the tumor as BVE with 54% probability, despite the case being BVW. The heatmap showed misplaced attention, with high focus on non-tumor regions (Figure 3e-i) and minimal attention (blue) on actual tumor areas (Figure 3e-ii). This inversion suggests that the model may have been confounded by dark staining and unclear boundaries that made it difficult to distinguish morphologically relevant features. Molecular profiling revealed an *NRAS* Q61R mutation, which may have contributed to mild cytologic activity, but not the classic features of *BRAF*-mutant melanoma (B et al. 2011). The incorrect spatial focus likely drove the misclassification in this case.

### Interpretation Category 3: BVE correctly predicted as BVE

In Figure 4a (Case 12), the model confidently predicted BVE status with 92% probability, accurately classifying the case. The heatmap showed strong attention over tumor regions, with most areas marked red. This alignment suggests the model correctly identified characteristic features of *BRAF*-mutant melanoma. A few pale tumor areas displayed slightly mixed attention, which may have contributed to a minor reduction in prediction confidence. Molecular data confirmed the presence of a *BRAF* V600E mutation with no concurrent *NRAS* or *NF1* mutations, supporting the model’s high-confidence output.

**Figure 4:**
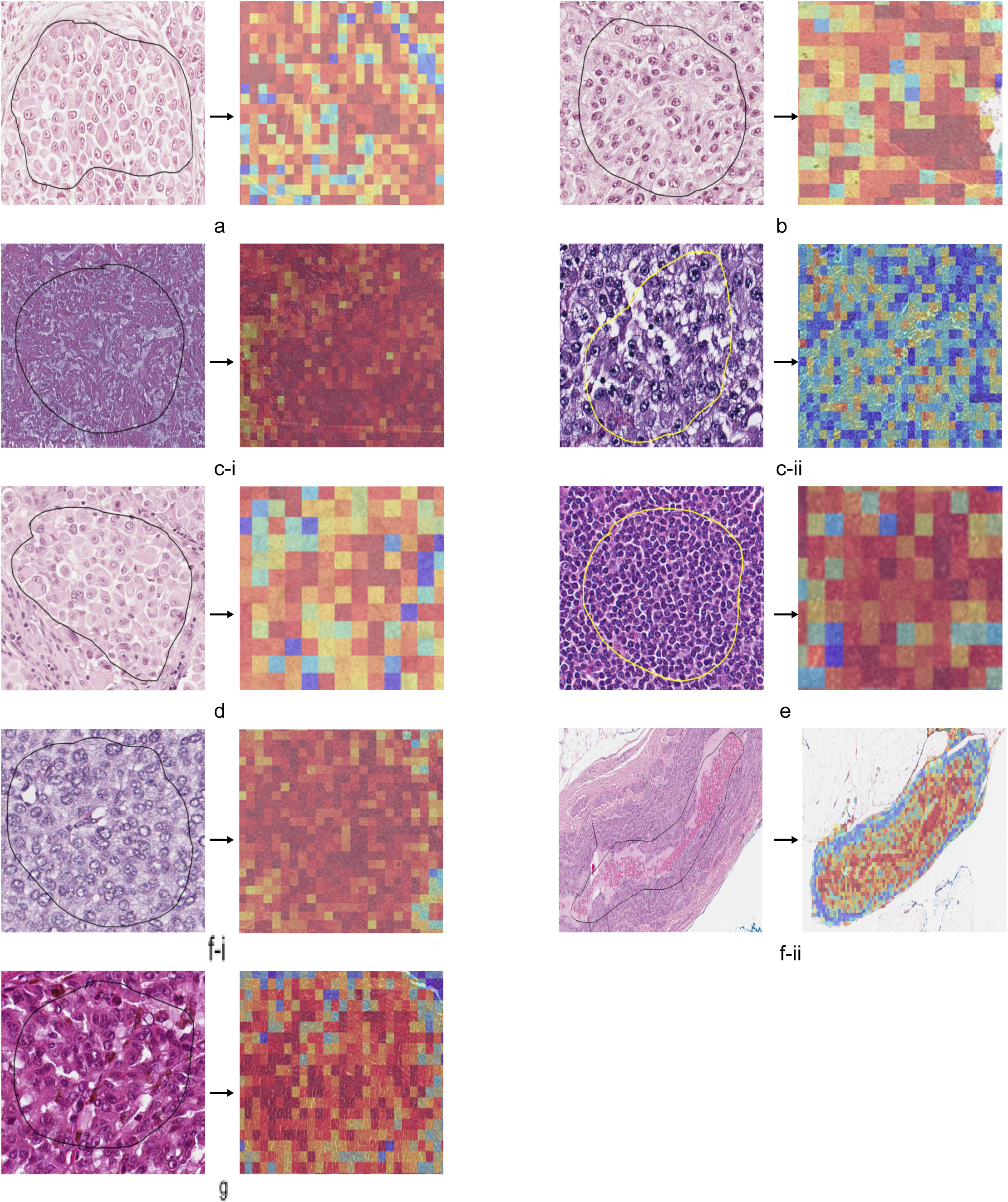
Representative Cases from Interpretation Category 3: BVE mutant correctly predicted as BVE. **(a)** Case 12 – TCGA-D3-A2JN: Heatmap correctly said 92% probability for BVE and 8% probability for BVW. ROI outlined in black,**(b)** Case 14 – TCGA-D3-A1Q9: Heatmap correctly predicted 89.7% probability for BVE and 10.3% probability for BVW. ROI outlined in black,**(c-i)** Case 15 – TCGA-ER-A2ND: Heatmap correctly predicted 86.1% probability for BVE and 13.9% probability for BVW; heatmap shows high attention over non-tumor areas. ROI outlined in black,**(c-ii)** Case 15 – TCGA-ER-A2ND: Heatmap correctly predicted 86.1% probability for BVE and 13.9% probability for BVW: heatmap shows low attention over tumor areas. ROI outlined in yellow,**(d)** Case 16 – TCGA-D3-A2J7: Heatmap correctly predicted 83.4% probability for BVE and 16.6% probability for BVW. ROI outlined in black,**(e)** Case 17 – TCGA-EE-A2MH:Heatmap correctly predicted 77.5% probability for BVE and 22.5% probability for BVW. ROI outlined in yellow,**(f-i)** Case 18 – TCGA-EE-A2GS: Heatmap correctly predicted 58.9% probability for BVE and 41.1% probability for BVW; heatmap shows attention over tumor areas. ROI outlined in black,**(f-ii)** Heatmap correctly predicted 58.9% probability for BVE and 41.1% probability for BVW; heatmap shows attention over non-tumor areas. ROI outlined in black,**(g)** Case 19 – TCGA-EE-A3AF: Heatmap correctly predicted 75.4% probability for BVE and 24.6% probability for BVW. ROI outlined in black.

In Figure 4b (Case 14), the model correctly predicted BVE status with 89.7% probability, focusing attention on the tumor regions, most of which appeared in red on the heatmap. While the slide exhibited typical *BRAF*-mutant morphology, the presence of scattered non-tumor cells may have introduced subtle confusion, slightly lowering the model’s prediction confidence despite the accurate classification.

In Figure 4c (Case 15), the model correctly predicted BVE status with 86.1% probability, even though the heatmap showed low attention (blue) over tumor regions (Figure 4c-ii) and high attention (red) over non-tumor areas (Figure 4c-i). This counterintuitive outcome suggests that the model may have focused on background features such as pigmented stroma or dense inflammatory zones that visually resemble BRAF-mutant histology in the training data. Although the actual tumor was not prioritized, these contextual similarities may have indirectly contributed to the correct prediction.

In Figure 4d (Case 16), the model correctly predicted BVE status with 83.4% probability, assigning strong attention (red) to tumor regions. However, the tumor cells appeared interspersed among normal cells, creating a scattered distribution. This spatial separation may have slightly reduced the model’s confidence, as the dispersed pattern could weaken the prominence of *BRAF*-mutant features and introduce noise from surrounding non-tumor tissue.

In Figure 4e (Case 17), the model correctly predicted BVE status with 77.5% probability, showing strong attention (red) across tumor regions. However, the slide was stained very darkly, which may have led the model to misidentify fibroblasts or other stromal elements as tumor cells. This misattribution could explain the slightly reduced probability, as non-specific features may have diluted the distinct morphologic signals typically associated with *BRAF*-mutant melanoma.

In Figure 4f (Case 18), the model correctly predicted BVE status with 58.9% probability, though the relatively close split suggests some uncertainty. The heatmap showed strong attention (red) over most tumor cells (Figure 4f-i) but also highlighted non-tumor regions (Figure 4f-ii), which may have introduced noise into the prediction. This overlap in attention could have confused the model, reducing its confidence despite the presence of clear *BRAF*-mutant morphology.

In Figure 4g (Case 19), the model correctly predicted BVE status with 75.4% probability. Most tumor regions were highlighted in red, indicating strong attention; however, some tumor areas were blue and the slide was notably dark-stained with melanin interspersed throughout. This combination of factors may have diluted the model’s confidence and reduced the predicted probability, despite the correct classification.

### Interpretation Category 4: BVE mutant incorrectly predicted as BVW

In Figure 5a (Case 13), the model incorrectly classified a BVE case as BVW with 97% confidence, assigning only 3% probability to the correct BVE label. Although the heatmap showed strong attention over tumor regions, the disorganized architecture, marked by cells oriented in multiple directions and lacking hallmark *BRAF*-mutant features, may have led the model to focus instead on more uniform, spindle-shaped areas typically associated with *BRAF* wild-type melanomas, reflecting a potential bias toward structural organization over cytologic detail (G et al. 2023). This misclassification highlights a key limitation in the current model’s ability to distinguish *BRAF*-driven tumors when the histologic presentation deviates from canonical mutant features. Notably, this was the only BVE case in the test set where such a confident misclassification occurred.

**Figure 5:**
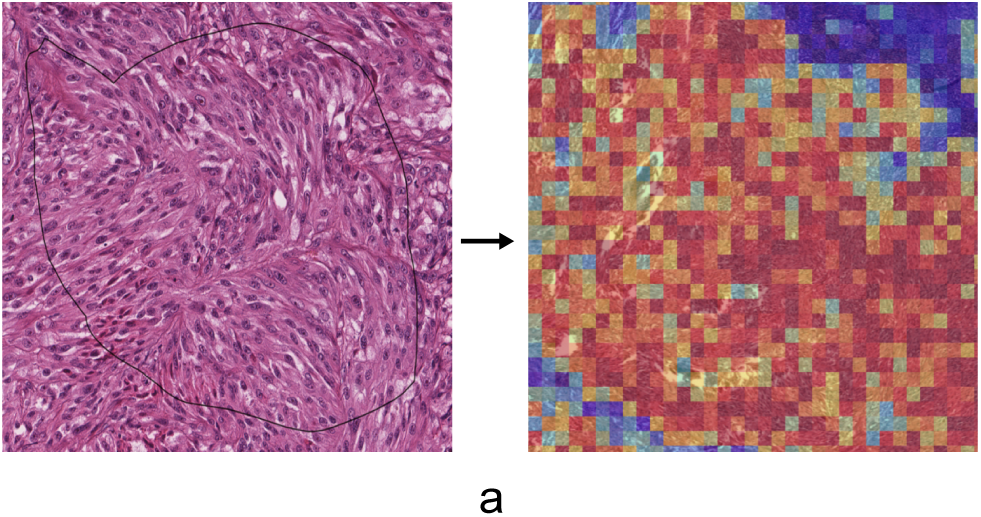
Representative Case from Interpretation Category 4: BVE mutant incorrectly predicted as BVW. **(a)** Heatmap falsely favored BVW with 97% probability, assigning only 3% probability to the correct BVE class. ROI outlined in black.

## DISCUSSION

This study presents XpressO-melanoma, an explainable deep learning model, capable of predicting *BRAF* V600E mutation status from diagnostic whole-slide images (WSIs) of cutaneous melanoma. The model achieved an AUC of 0.79 and a precision and recall rate of 70% on an independent test set, demonstrating its potential to detect morphologic correlates of *BRAF* mutations using only standard histopathology. In addition, the model exhibited a positive prediction rate of 87.5% for *BRAF* V600E-mutated cases (BVE) and 54.54% for *BRAF* V600E wild-type cases (BVW). These rates reflect the model’s relative sensitivity to the two mutation states, further highlighting its utility in identifying histopathologic features associated with *BRAF* mutation status. Importantly, the attention-based architecture enabled spatial localization of tumor regions most predictive of mutation status, enhancing the transparency and interpretability of model predictions.

In interpreting model predictions, we observed strong agreement between high-attention regions and known morphologic differences between BVE and BVW melanomas. Cases predicted as BVE often showed epithelioid morphology, prominent nucleoli, cytoplasmic pigmentation, and multinucleation which are features commonly associated with *BRAF* V600E-mutant melanomas (Lee et al. 2018). On the other hand, cases predicted as BVW showed ovoid nuclei, dense fibrous stroma, and lower degrees of pleomorphism. These align with descriptions of *BRAF*-wild-type subtypes, including desmoplastic and spindle-cell melanomas (Se et al. 2014). This pattern suggests that the model successfully learned to associate relevant histologic features with mutation status.

Interestingly, even in cases where the model misclassified a wild-type tumor as mutant, the histology often reflected underlying oncogenic drivers other than *BRAF*. Among the 11 BVW-labeled cases, several carried *NRAS* or *NF1* mutations (e.g., *NRAS* Q61K/L/R; *NF1* R440*, C383Yfs*), which are known to drive intermediate morphologies, such as increased cellularity or desmoplastic features, that partially overlap with *BRAF*-mutant characteristics. Only two misclassified wild-type cases were truly triple-wildtype. This suggests that the model’s attention was not entirely misplaced, but rather sensitive to broader oncogenic patterns. Notably, there was only one instance in which a BRAF-mutant case was predicted as BRAF wild-type. The model’s ability to rarely miss BRAF-mutant cases highlights its potential to support clinically meaningful diagnostic decisions. Overall, such predictive flexibility is especially valuable in resource-limited settings, where confirmatory molecular profiling may not be available, and histology may be the only source of diagnostic insight.

At the same time, we also encountered borderline cases where the morphology was more ambiguous. In such examples, the model often produced nearly equal probabilities (50-50) for both classes or made incorrect predictions. These occurred in slides showing intermixed spindle cells, necrotic cells, low tumor cellularity, or prominent reactive stroma - patterns known to complicate histopathologic interpretations (Colombino et al. 2024). The overlap in features between *BRAF* V600E mutant and *BRAF* V600E wild-type melanomas can make interpretation difficult, especially in fibrotic or inflamed areas. The model’s behavior in these challenging cases indicates sensitivity to subtle or context-dependent features. This highlights the importance of incorporating more refined morphologic descriptors in future work.

Notably, the current model was trained without any manual annotation or human-directed labeling of tumor regions. It relied on weak supervision using only slide-level mutation labels. Despite this, the model consistently focused on histologically meaningful regions, indicating its ability to learn features associated with BRAF mutation status. Although this strategy offers promising scalability, it does not fully capture the complexity of tumor morphology on its own. In a few instances, such as cases where attention extended to stromal or necrotic regions - the model highlighted areas that may not be directly informative for mutation classification. These observations suggest that incorporating expert annotation could further guide and refine the model’s spatial focus.

To this end, our upcoming work will integrate pathologist-guided annotation of melanoma tumor regions alongside computational feature extraction. By combining deep learning with curated morphologic input, we aim to capture richer phenotypic detail, clarify ambiguous predictions, and reduce false positives. This hybrid approach is expected to sharpen the model’s sensitivity to BRAF-related histologic patterns and improve spatial reliability in clinical contexts (Kim et al. 2022).

Additionally, although the current model is designed to distinguish only between BRAF V600E-mutant and wild-type classes, its sensitivity to NRAS- and NF1-driven morphologies indicates that it has implicitly learned features reflective of broader oncogenic phenotypes. This is evident in several BVW cases with NRAS or NF1 mutations that were predicted as mutant, highlighting the model’s ability to detect intermediate histologic cues shared across mutation subtypes. Building on this, our future work will extend the model to a multi-class framework capable of distinguishing BRAF, NRAS, NF1, and triple-wildtype cases. This evolution will support more granular mutation prediction and increase the model’s clinical relevance across diverse melanoma subtypes.

Our pipeline demonstrates a scalable, low-cost method to infer clinically actionable molecular information directly from routinely collected WSIs. Such a tool could be particularly valuable in resource-limited healthcare settings where diagnostic molecular testing is limited, or in cases where biopsy material is insufficient for additional genomic assays. By integrating AI into the diagnostic workflow, pathologists could be equipped with early mutation probability scores to prioritize confirmatory testing or guide treatment discussions. Nevertheless, thoughtful interpretation of these scores remains important, as the model may occasionally prioritize regions that are histologically ambiguous. In future phases, we will incorporate expert annotations to guide and refine model predictions.

In conclusion, we present XpressO-melanoma, an attention-based deep learning model capable of predicting BRAF V600E mutation status from melanoma WSIs with high interpretability and promising predictive performance. By integrating pathomic features and expanding dataset diversity, the model’s predictive capacity and clinical utility can be further optimized, setting the foundation for multi-modal diagnostic frameworks in computational pathology.

## MATERIALS AND METHODS

### Collection and Processing of WSIs

Diagnostic (Dx) Whole-Slide Images (WSIs) of cutaneous melanoma were obtained from The Cancer Genome Atlas (TCGA) portal using the Mutation Frequency tool (The Cancer Genome Atlas Program, National Cancer Institute, 2025). A total of 192 cases were collected, comprising 112 *BRAF* V600E wild-type (BVW) tumors and 80 *BRAF* V600E mutant (BVE) tumors, as defined by their DNA mutation profiles available through TCGA.

To facilitate computational analysis, the WSIs were processed using the XpressO pipeline (Sukhadia et al. 2025), an explainable deep learning (DL) framework initially developed for weakly supervised gene expression prediction in invasive breast cancer using TCGA datasets. The same modular pipeline was subsequently adapted for the analysis of cutaneous melanoma WSIs in this study, except that the mutation status labels were used instead of gene expression labels. XpressO integrates several key components to enable end-to-end histopathological analysis.

As part of XpressO, the Clustering-constrained Attention Multiple Instance Learning (CLAM) module (My et al.) was used to segment tumor regions of interest (ROIs) on each WSI. CLAM leverages an attention mechanism to automatically identify histologically significant tumor areas without requiring manual annotations, thereby improving both model interpretability and classification accuracy.

Given the high resolution of WSIs, XpressO’s OpenCV-based patching module employs a sliding window approach to divide each tissue section into manageable, non-overlapping patches of 256×256 pixels at 20x magnification (Python Software Foundation,2025). Non-informative patches, such as blank regions or heavily stained areas, are filtered out to retain only viable tumor tissue.

For feature extraction, XpressO utilizes the UNI module (Unified Network for Instance-level Representation Learning) (J et al.), a pre-trained, weakly supervised Vision Transformer (ViT-L/16) architecture optimized for histopathological image analysis (Dosovitskiy et al. 2021; Oquab et al. 2024). Each retained patch is encoded into a high-dimensional feature embedding, yielding a compact and standardized representation of the tissue’s morphological features. These embeddings are subsequently used for downstream classification tasks.

### Dataset Splitting and Label Assignment

The 192 cutaneous melanoma WSIs used in this study were assigned binary labels (0 or 1) based on their *BRAF* V600E mutation status derived from TCGA survival plots. Each TCGA case ID was mapped to a binary label based on the associated mutation class: cases labeled as “BVW” were classified as *BRAF* V600E wild-type (label = 0; n = 112), and those labeled as “BVE” were designated as *BRAF* V600E mutant (label = 1; n = 80). This labeling scheme was derived using a survival-based mutation stratification of TCGA melanoma cases. These labels were applied at the slide (case) level and were used for supervised learning. Since the classification task involved a binary endpoint, no additional transformation or thresholding was necessary.

### Model Training

The analysis pipeline began with the segmentation of whole slide images (WSIs) using the Segmentation module of the XpressO framework to segment the tumor regions of interest (ROIs) using the CLAM model (My et al.; Sukhadia et al. 2025). Following segmentation, patch-level features were extracted from the identified regions of interest (ROIs) using XpressO’s UNI model, which is a pre-trained foundation model capable of identifying multiple tumor types including melanoma (J et al.; Sukhadia et al. 2025). The extracted feature embeddings were then utilized for classification of mutation status using the CLAM SB model (My et al.; Sukhadia et al. 2025). This weakly supervised learning framework aggregated patch-level information without requiring pixel-level annotations, allowing the model to learn discriminative features that contribute to slide-level *BRAF* mutation status.

Further, an attention-based pooling layer was applied during training to assign weights to individual patch embeddings, determining their relative importance for predicting the slide-level *BRAF* mutation status. Higher attention scores indicated regions with greater influence on the mutation classification outcome (i.e., *BRAF* V600E wildtype versus *BRAF* V600E mutant). The weighted patch embeddings were then aggregated into a single slide-level representation, which was used to generate the final classification output for *BRAF* mutation status.

Within the XpressO pipeline, training was performed using the Adam optimizer with a learning rate of 2×10, a batch size of one WSI per iteration, and dropout set at 25% to reduce overfitting. The model was trained for 200 epochs with early stopping triggered if validation loss failed to improve for 10 consecutive epochs. The cross-validation fold ranged from 0 to 11.

### Model Evaluation and Metrics

The complete dataset of 192 WSIs got partitioned randomly at the patient level into three independent sets: 154 cases (80%) were used for training the model, 19 cases (10%) for validation (hyperparameter tuning and early stopping), and 19 cases (10%) were reserved as a fully independent test set. This partition corresponded to the best performing fold selected during cross-validation. The label distribution across these splits showcased a balanced representation of both BVE and BVW classes (Figure 1).

Model performance was assessed using the Evaluation module within the XpressO pipeline, which is designed to support standard binary classification tasks in histopathological analysis. This module leverages Python’s scikit-learn library (Python Software Foundation,2025; Sukhadia et al. 2025) to compute key performance metrics, including the area under the receiver operating characteristic curve (AUC-ROC), accuracy, precision (positive predictive value), recall (sensitivity), and F1-score (the harmonic mean of precision and recall). Among these, the final AUC-ROC of the Test set was used as the primary indicator of the model’s discriminative capacity to distinguish between BVE and BVW melanomas, while the remaining metrics provided insight into predictive reliability in a binary classification context.

### Attention Heatmap Generation and Visualization

Attention heatmaps were generated on the test set using the Heatmap module from XpressO (Sukhadia et al. 2025). For each WSI, the model’s attention scores assigned to individual patches were mapped back onto their corresponding spatial coordinates. Patches with higher attention weights were visualized with greater intensity (red), enabling the identification of regions that contributed most strongly to the slide-level *BRAF* mutation prediction. Attention scores representing the probability of mutation-related features were aggregated across the top-10 ranked patches, which were then visualized. These scores were overlaid onto the original WSIs to highlight tumor regions of high diagnostic relevance and were inspected further for their truthiness by an experienced pathologist.

## AUTHOR CONTRIBUTIONS STATEMENT

Conceptualization TP, SS; Data curation VR; Formal analysis VR; Funding acquisition SS (self); Investigation VR TP SS; Methodology VR SS; Project administration SS; Resources SS; Software SS VR; Supervision TP, SS; Validation VR, TP, SS; Visualization VR, TP, SS; Writing – original draft VR; Writing – review & editing VR, TP, SS

## Abbreviations Used

AI: Artificial Intelligence
AUC: Area Under the Curve
BRAF V600E: B-Raf Proto-Oncogene, Serine/Threonine Kinase, Valine 600 Glutamic acid mutation
BVE: BRAF V600E mutant
BVW: BRAF V600E wild-type
CLAM: Clustering-constrained Attention Multiple Instance Learning
CNN: Convolutional Neural Network
DL: Deep Learning
Dx: Diagnostic
H&E: Hematoxylin and Eosin
IHC: Immunohistochemistry
IF: Immunofluorescence
MIL: Multiple Instance Learning
NGS: Next-Generation Sequencing
NF1: Neurofibromin 1
NRAS: Neuroblastoma RAS Viral Oncogene Homolog
PCR: Polymerase Chain Reaction
ROI: Region of Interest
RT-PCR: Real-Time Polymerase Chain Reaction
TCGA: The Cancer Genome Atlas
UNI: Unified Network for Instance-level Representation Learning
ViT: Vision Transformer
WSI: Whole Slide Image

